# Cryo-EM structures reveal the activation and substrate recognition mechanism of human enteropeptidase

**DOI:** 10.1101/2022.03.07.483351

**Authors:** Xiaoli Yang, Zhanyu Ding, Lisi Peng, Qiuyue Song, Fang Cui, Deyu Zhang, Chuanchao Xia, Keliang Li, Hua Yin, Shiyu Li, Zhaoshen Li, Haojie Huang

## Abstract

The enteropeptidase (EP) initiates the intestinal digestion by proteolytic processing of trypsinogen, generating catalytic active trypsin. The dysfunction of EP will cause a series of pancreatic diseases, the most severe of which is acute necrotizing pancreatitis. However, the molecular mechanism of EP activation and substrate recognition remain elusive due to the lack of structural information, hampering the structure-based research of EP and even further EP-targeted drug design. Here we report cryo-EM structures of human EP in multiple states, covering the functional cycle spanning from inactive to active state and eventually to the substrate binding state, with the inactive core region reached an atomic 2.7-Å-resolution. The heavy chain of EP exhibits a clamping configuration with the CUB2 domain serving for substrate recognition. The N-terminus of light chain induces the surface loop remodeling from inactive to active conformation, resulting in a highly dynamic and active EP. Then the heavy chain performs like a hinge to ensure the flexibility of light chain for substrate recruitment and subsequent cleavage. Our study provides structural insights of EP remodeling and heavy chain dynamics while performing enzymatic function, facilitating our understanding of the pathogenies of EP-related pancreatitis and the EP-targeted treatment of pancreatitis.

## Introduction

Enteropeptidase (EP), also named enterokinase, localized to the brush border of the duodenal and jejunal mucosa ^1, 2^, is an essential enzyme in food digestion and the physiological activator of trypsinogen ^2, 3^. Activated EP could stimulate the conversion of pancreatic trypsinogen to trypsin, via cleavage of the specific trypsinogen activated peptide (Asp-Asp-Asp-Asp-Lys, DDDDK), which is highly conserved in vertebrates ^4-6^. Hereafter, trypsin initiates a cascade of activation of various pancreatic zymogens, such as chymotrypsinogen, proelastase, procarboxypeptidase and prolipase, to regulate nutrient absorption and metabolism ^2, 7-10^. The dysfunction of EP will cause a series of diseases. Newborns with congenital deficiency of EP will need pancreatic enzyme replacement therapy or be treated with a mixture of amino acids in the diet to promote growth and normal health, suggesting the pivotal role of EP in regulating body homeostasis ^1, 11-14^. On the other hand, EP redundancy will also cause diseases. Combined caerulein with EP infusions resulted in the transformation of mild to necrotizing pancreatitis in rat models ^15^. Furthermore, duodenopancreatic reflux of activated EP can trigger the pancreatic enzyme cascade, leading to acute necrotizing pancreatitis ^2, 16-19^. Additionally, translocation of EP into the pancreatic-biliary tract has been shown to associated with pancreatitis ^20^. Therefore, EP has attracted intense interests as a drug target ^10, 21, 22^.

EP is a type II transmembrane serine protease, synthesized as a zymogen without catalytic activity, until activated by trypsin or related proteases ^6, 23, 24^. EP contains a multidomain heavy chain and a catalytic light chain, linked by a disulfide bond ^25^. The physiological activator of EP and the mechanism of EP activation remain unestablished ^24, 26^. The heavy chain contains a single-pass transmembrane (TM) domain and seven structure motifs, including one copy of SEA, MAM, SRCR, and two copies of LDLR, CUB ^25, 27^. The function of these domain and motifs are predicted to involve in protein anchoring, macromolecular substrate recognition and inhibitor specificity ^24, 25, 28, 29^, while the exact roles still need further exploration. The light chain is homologous to trypsin-like serine proteinases with a typical Aspartate (D) - Histidine (H) - Serine (S) catalytic triad, responsible for peptidase activity ^10, 28^.

Structural exploration of full-length EP is crucial to understand how heavy chain modulate the fully function of EP. Existing crystal structures of EP contains only the light chain, showing a typical α/β trypsin-like serine protease fold ^10, 30-32^. One EP structure with bound inhibitor elucidates the inhibition mechanism of EP light chain by camostat ^10^. However, the inhibitor specificity of EP mainly depends on the heavy chain ^24^. Another structure of EP light chain in complex with the substrate peptide was also solved, however, the proteolytic activity toward trypsinogen is significantly impaired with deletion of the heavy chain ^2, 4, 6, 30, 33^. Consequently, there is still a huge gap in the understanding of EP functional mechanism due to the lack of structural information on heavy chain. The structural analysis of full-length EP and its complex with physiological substrate will be an important step to outline a framework for detailed EP functionality characterization, and serve as the foundation for EP-targeted drug design.

Here we determine cryo-electron microscopy (Cryo-EM) structures of multiple conformations of human EP (hEP) from inactive to active states, and in complex with nafamostat and trypsinogen. Our results reveal the architecture of full-length hEP and provide a mechanistic model for hEP catalytic function, which advances our understanding of the mechanism of EP dynamics and substrate coordination, and provides structural basis for the targeted therapy of related diseases.

## Results

### Inactive state of human enteropeptidase (hEP)

We expressed hEP (182-1,019 residues) in HEK293F cells (Fig. 1A). The purified hEP is a single-chain zymogen with a large molecular weight (>130 KDa), much higher than its theoretical molecular weight of 95 KDa, indicating our hEP is heavily glycosylated ^24^ (Fig. S1A). Our in vitro enzymatic assay confirms this hEP is inactive, by showing no detectable activity toward substrate cleavage, which is consistent with previous reports ^2, 24^ (Fig. S1B). When the as-produced hEP was cleaved into two chains by trypsin, its protease activity is completely shown comparable to commercial active one (Fig. S1A-B). This hEP was subjected to cryo-EM single particle analysis, enabling us to capture the inactive state of EP to 3.8 Å resolution, with the core region reached an atomic 2.7-Å-resolution (Figs. 1B, S1C-H, and Table 1). The structure is observed to be ∼130 Å in length, ∼50 Å in width, and ∼80 Å in height (Fig. 1B). This is the first experimental structure of EP containing heavy chain, revealing the overall shape how the heavy chain clamps the light chain (Fig. 1B).

**Fig. 1.**
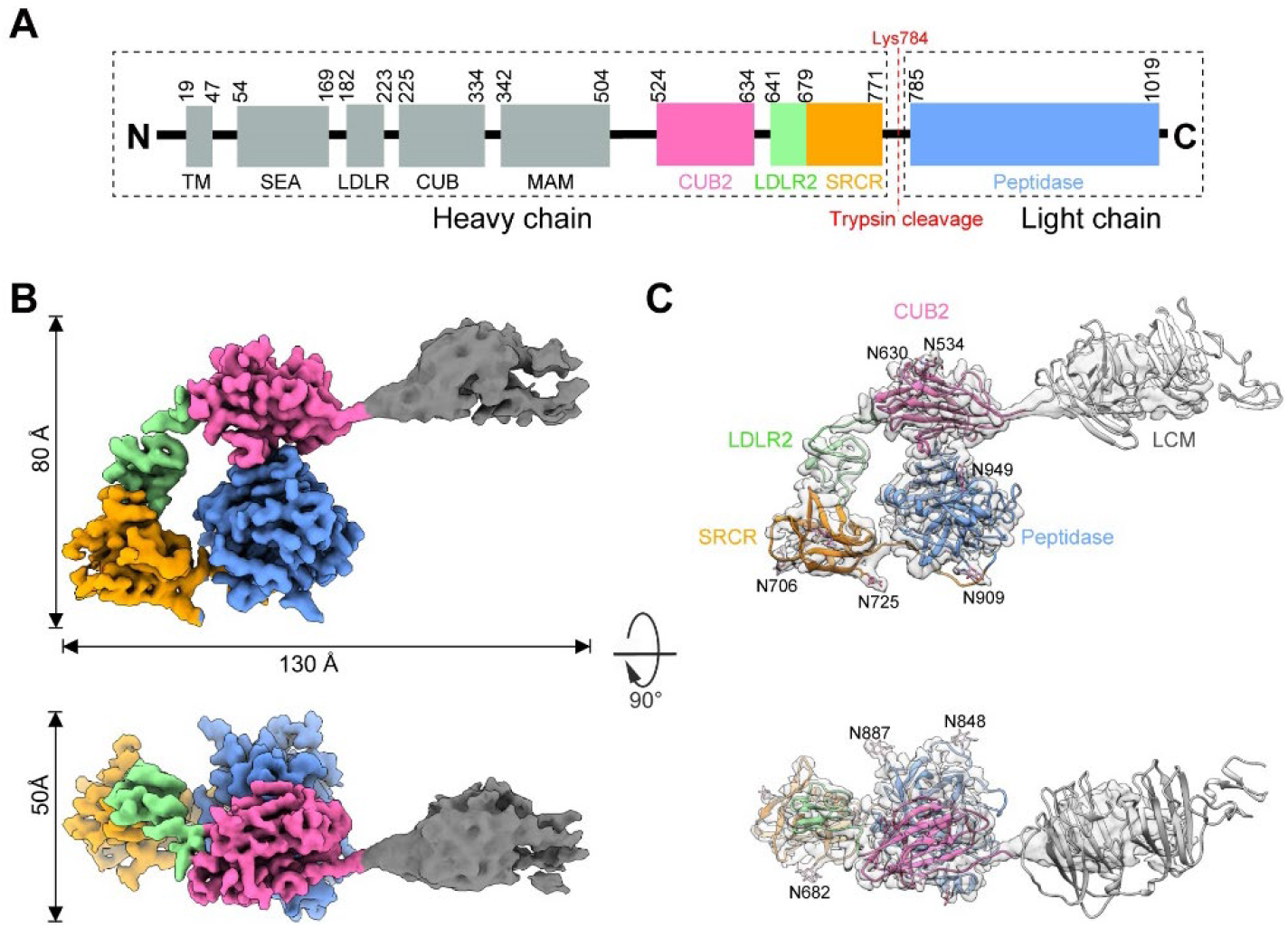
Cryo-EM structure of hEP in the inactive state. **(A)** Domain information of hEP. Numbers indicate the position of each domain in the amino acid sequence. **(B)** Cryo-EM structure of inactive EP. The colored density map is a composite map generated with locally refined EP-core and the low-resolution region in the whole map. The domain color scheme is followed in **(A)**. Unless otherwise indicated, the same domain color scheme is applied to all figures. **(C)** The structure model of the inactive EP, with the map in transparent. The detected 9 N-linked glycans were indicated by atomic view and the covalent attached amino acids were also labeled.

The peptidase domain is well resolved with the amino acid side chains clearly detected, except the surface loops (Fig. S2A-B). These loops, like those of other serine proteases in the inactive state, mainly L1, L2, and LD ^34, 35^, are severely disordered having no density to fit in (Fig. S2A). For the heavy chain, the SRCR, LDLR2, and CUB2 domains were unambiguously build into the density map (Fig. 1C). The SRCR and CUB2 domains grab the light chain increasing the interaction interface to 1163.2 Å^2^ (Fig. S2C). The map displays the orientation of the remaining three domains (LDLR, CUB and MAM, LCM for short) with a few details (Fig. 1B-C). These domains show fuzzy density without structural features from the 2D class averages, suggesting their mobility relative to the core region (Fig. S1E). To further confirm the orientation of these domains, we searched the AlphaFold Protein Structure Database for the full-length EP model (Jumper et al., 2021; Tunyasuvunakool et al., 2021). The predicted full-length structure depicts a scenario that the light chain is deeply enclosed by heavy chain in EP precursor (Fig. S2D). Fitting the predicted model into our inactive EP map, the core domain matches perfectly with our electron density, and the RMSD between the predicted model and the experimental one is 0.935, demonstrating the reliability for the 3D structure predication of AlphaFold2 (Fig. S2D). The predicted LCM domains extend in almost the same direction as the density we have resolved, except for a slightly shift of ∼20 Å (Fig. S2E).

EP is a highly glycosylated protein with 18 potential N-linked glycosylation sites, of which 9 glycans located in the core region (Fig. S2F). The removal of glycosylation may change its tertiary structure and enzymatic activity ^25^. Our SDS-page indicated the EP is heavily glycosylated (Fig. S1A), indeed we detected all the 9 N-linked glycans in our inactive EP core region, with the density clearly resolved (Fig. S2G). This suggested our structure was in a nearly physiological state, suitable for the exploration of EP functionality.

### Activation mechanism of hEP

Trypsin cleaves the EP into heavy and light chains to fully activate its catalytic ability. Two chains appearing as two bands on our SDS PAGE and the high protease activity indicate trypsin-treated EP is indeed in the active state (Fig. S1A-B). Though these two bands containing the complete length of expressed residues, only the core region density could be solved. We performed deep classification as that for inactive dataset, still, we could not find any trace for the previously low-resolution LCM domains (Fig. S3C and S3F). It suggested that these unresolved LCM domains are much more dynamic relative to the core region in the active state. Thus, we only solved this activated EP-core cryo-EM structure to an overall resolution of 3.2 Å, allowing us to further explore the activation mechanism of hEP (Figs. 2A, S3, and Table 1). The overall shape of the activated EP-core is quite similar to that of the inactive core map, with the heavy chain clamps the catalytic light chain (Fig. 2A and S4A). However, superimposing of these two core maps exhibit more density near the catalytic site in the active map (Fig. S4A). These extra densities corresponding to the loops, L1, L2, and LD (Fig. 2B), which were not resolved in inactive EP (Fig. S2A). Overlaying our activated structure with the published crystalized model (PDB ID: 4DGJ) shows overall a good match, with the catalytic pocket well determined by surface loops, the connected three β-strands, and the disulfide bond Cys967-Cys995 (Figs. 2B-C and S4B). This pocket organization is attributed by the newly-formed N-terminus of the light chain after cleavage, which is the conserved N-terminal activation peptide sequence (IVGG)^36^. This sequence turns over ∼180 degrees, and Ile785 shifts by 16.6 Å to probably stabilize loop L1 by a salt bridge with the side chain of Asp970, allosterically steady the catalytic center of EP compared to that in the inactive state (Fig. 2C-E).

**Fig. 2.**
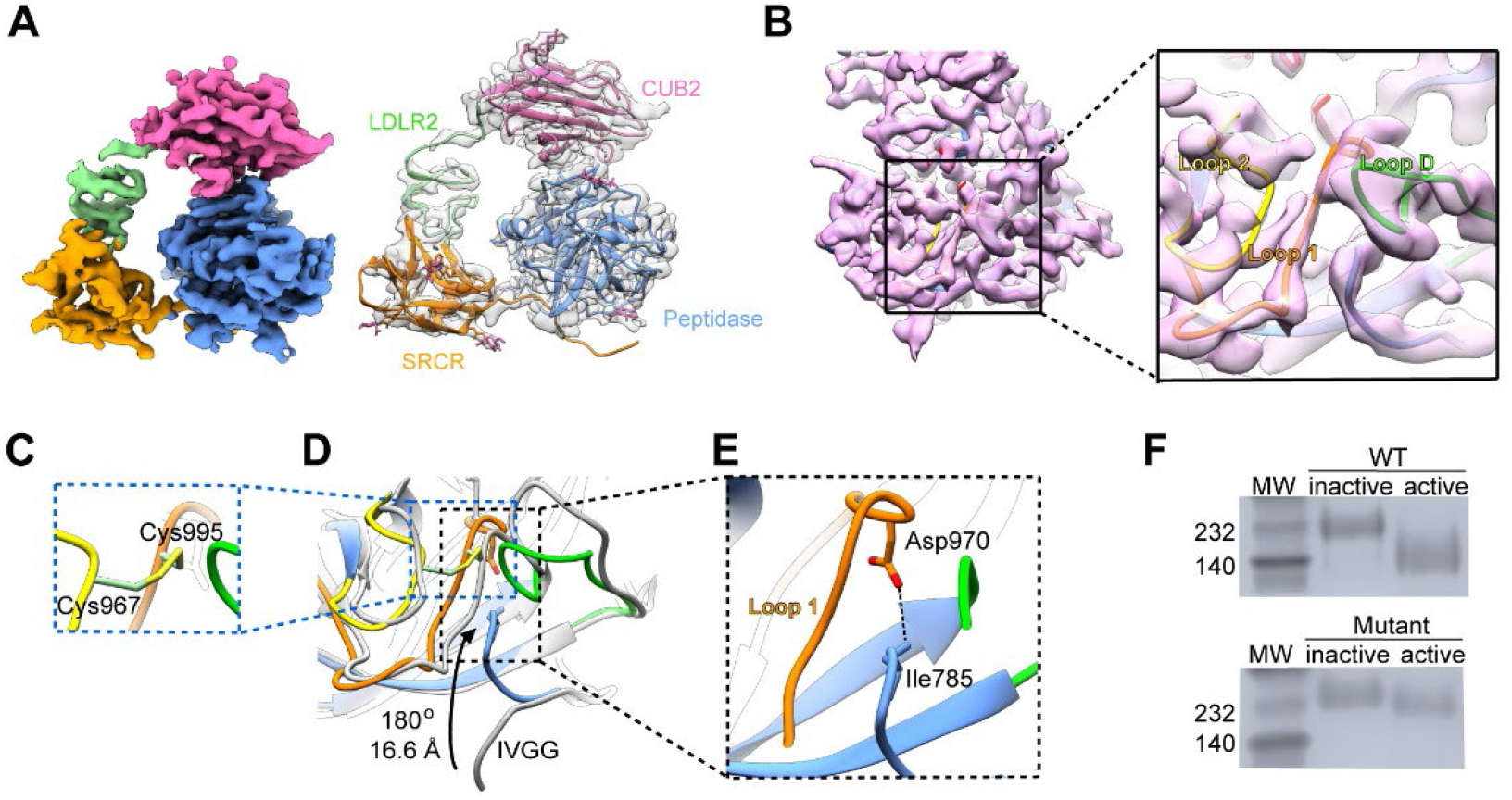
Activation mechanism of hEP. **(A)** Cryo-EM structure of activated wtEP-core, with model fitted in. The detected 9 N-linked glycans were indicated by atomic view. **(B)** The peptidase domain of activated wtEP-core map fitted with the corresponding pseudo-atomic model, with the surface loops enlarged for clarity. **(C)** The interdomain disulfide pair (Cys967-Cys995) forms to serve as the wall of the catalytic pocket. **(D)** Structural comparisons between active (with different colors) and inactive (grey) states. The black arrow indicates the rotation and shift of the IVGG sequence. **(E)** The Asp970:Ile785 salt bridge stabilize the loop 1. **(F)** The activation of EP monitored by a gel-shift assay. The wtEP shows a supershift upon activation, while the shift minishes for the triple mutant of EP.

During the conformational changes of the IVGG sequence, the catalytic triad, D876-H825-S971, seems to accommodate a little conformational change (Fig. S4C). To test whether the catalytic triad has influence on the activation of EP, all three residues were mutated to alanine simultaneously, and the enzymatic assay confirmed this mutated EP loss of proteolytic activity (Fig. S1B). Noteworthy, our native gel-shift assays showed a super shift upon activation for wild-type EP, while the shift minishes for the triple mutant of EP (Fig. 2F), indicating the charge variations occurs.

To investigate the influence of charge variation on the catalytic pocket rearrangement, we determined this activated mutant EP structure with core region to an overall resolution of 3.7 Å (Figs. S3D-E\, S3G-I, S4D, and Table 1). Comparison of the two active state maps indicate they are almost identical (Fig. S4E-G). The mutated residues were confirmed by side chain density (Fig. S4H). Hence, the mutation of the catalytic triad abolishes the catalytic activity, but do not introduce any significant catalytic pocket rearrangement. Putting together, the enzymatic activity of EP is initiated by the flipping of IVGG to stabilize the L1, L2, and LD loops surrounding the catalytic sites. The surface charge provided by the catalytic triad is believed to have no effects on the EP activation, but could affect catalytic substrate reaction probably by lowering substrate binding affinity.

### Inhibition of hEP

Inhibition of the serine protease becomes a new therapy for the treatment of many diseases, including Alzheimer disease ^37^, autoimmunity disease ^38^, and even COVID-19 pandemic ^39, 40^. The development of specific inhibitor that selectively target EP maybe a strategy to the clinical intervention of pancreatitis ^10^. Different kinds of inhibitors were tested against inhibitory effects to hEP, among which, nafamostat and camostat showed the best performance (Fig. 3A). Nafamostat was found to be more effective than camostat in inhibiting other serine protease ^41, 42^, and was reported to be clinically useful in the therapy of acute pancreatitis ^43, 44^. Given that, nafamostat is a broad-spectrum synthetic serine protease inhibitor (Fig. 3C), with the IC50 value of 1.5 μM for the inhibition of EP activity, we choose nafamostat to investigate the inhibition mechanism of hEP. Not to mentioned that the crystal structure of EP with camostat is already available.

**Fig. 3.**
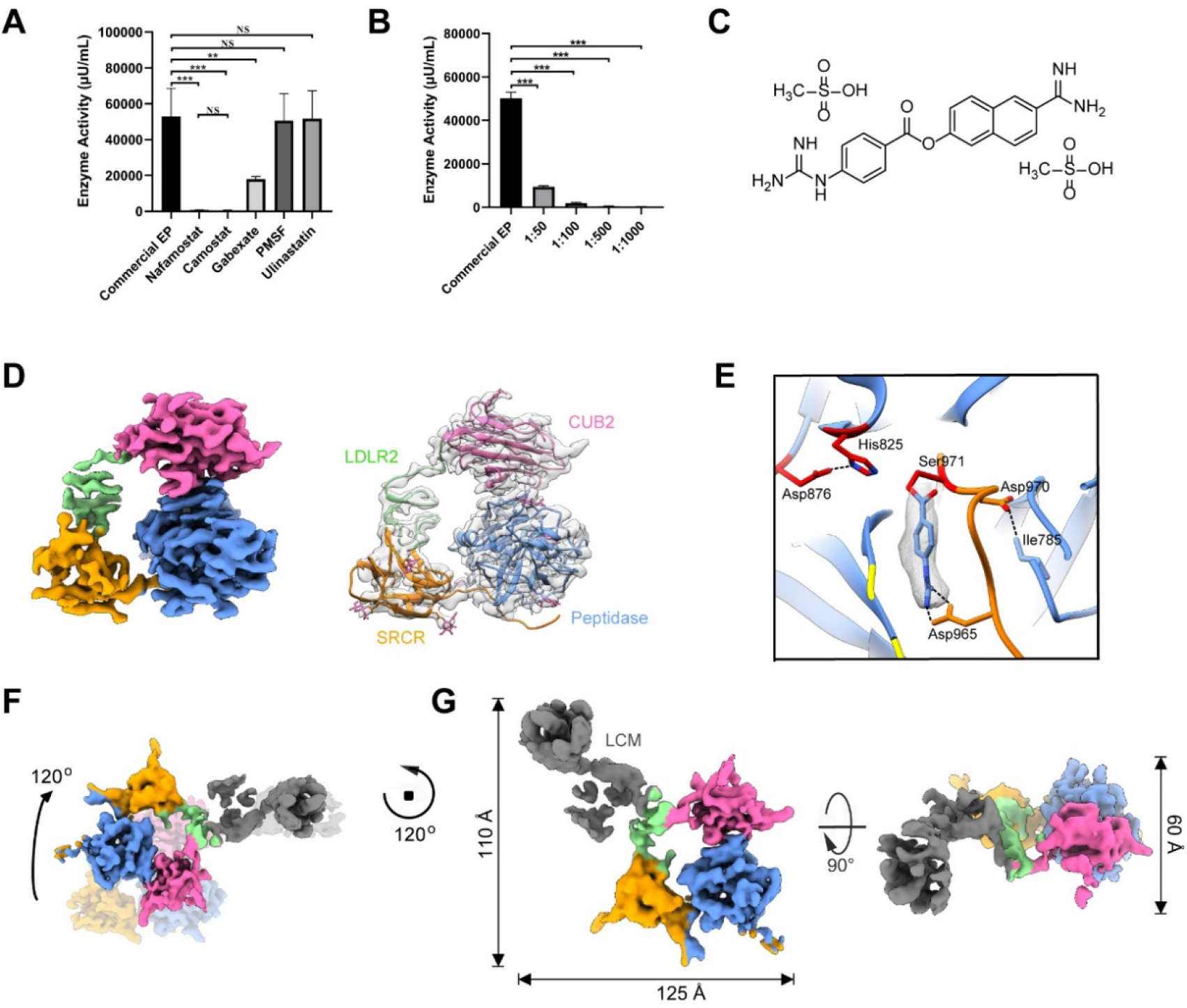
Structural features of inhibited hEP. **(A)** *In vitro* EP activity experiment under different inhibitor conditions. Commercial EP was also tested as a positive control. SDs (standard deviations) were calculated from at least three independent experiments. Significance was tested using one-way ANOVA; ^**^*P*<0.01, ^***^*P*<0.001, ^NS^*P*>0.05. **(B)** *In vitro* EP activity experiment. HEPs were incubated with different amount of nafamostat. Commercial EP was also tested as a positive control. SDs (standard deviations) were calculated from three independent experiments. **(C)** Chemical structure of nafamostat mesylate. **(D)** The cryo-EM structure of inhibited EP core, with model fitted in. The detected 9 N-linked glycans were indicated by atomic view. **(E)** Interactions between nafamostat and hEP. **(F)** Overlaying of the inactive EP (light shadow) with inhibited EP-complete maps illustrates the conformation change of the core region. **(G)** Cryo-EM structure of inhibited EP-complete map.

We incubated activated hEP with nafamostat at different molar ratios, and 1:1000 is chosen for structural determination due to the best inhibitory effect (Fig. 3B). Similar to the activated dataset, ∼75% good particles showed no density of LCM domains, resulting an EP-core map at a resolution of 3.1 Å (Figs. 3D, S5, and Table 1). Surprisingly, within the light chain domain, an unambiguous extra density was found in the catalytic pocket, which is assigned to the nafamostat adduct (Figs. 3E and S6C). The binding affinity was found to be as high as 9.51E-05, indicating nafamostat as a promising drug candidate (Fig. S6D). Nafamostat forms a covalent bond with the catalytic residue Ser971, and shows polar contact with the conserved Asp965 (Fig. 3E), consistent with that for camostat in inhibited EP light chain crystal structure (Sun et al., 2020). This blockage occupancy of catalytic residue excludes the binding of substrate, inhibiting the cleavage activity of EP. The binding environment of nafamostat in our structure is quite similar to that in other serine protease (Mahoney et al., 2021), explicitly verifying the inhibition mechanism of serine protease through covalent and occupancy (Sasaki et al., 2019). Still, we should mention this inhibited EP-core map is almost identical to that in active state (Fig. S6A-B).

Probably owing to the inhibitor binding, we captured the rest 25% good particles in a complete profile of EP, with the LCM domains density clearly shown (Figs. 3F-H, S5, and Table 1). The height of this inhibited EP-complete structure increases to ∼110 Å from ∼80 Å of inactive EP, resulting from the ∼120 degrees rotation between the EP-core and LCM domains (Fig. 3F). This dramatic flexibility is realized by the loop between MAM and CUB2, performing like a hinge. Consequently, the LDLR2 domain relocated to contact the LCM domains and further cause a change in the position of LDLR2 (Fig. S6F). The light chain remains firmly clamp-hold by a few domains of the heavy chain within the core region (Fig.S6E), forming a rigid integrity to assure the intimate interaction between heavy and light chains after the cleavage for EP activation.

### Structural characterization in different states

To measure the conformational mobility variances in different functional states of hEP, we calculated the B-factors for the well-solved models. Generally, the CUB2 and peptidase domains have the lowest B-factors in all structures involved, indicating the quiescent nature of these two domains (Fig. S7A). Comparing all the states, the overall B-factors of inactive and inhibited states were obviously lower than that of the active state (Fig. S7A), reminiscent that they are both deactivated. In another word, inactive EP favors the low energy to keep steady until the folded activation domain forms, being a high-energy activation state ready to act as a catalyst; the inhibitor bound to catalytic residue would reduce the global energy, returning the whole enzyme to a low energy state.

Surprisingly, the patterns of surface property around the catalytic pocket of inactive EP were observed to be different from those in other states (Fig. S7B), exhibiting negative charges. While in the activated state, the positive surface charges around the catalytic center probably confers specificity for recognize the negatively charged acidic residues of trypsinogen activation peptides (DDDDK) These findings, to some extent, would facilitate our understanding of the mechanism of EP functionality.

### Substrate-engaged EP

Given the critical role of pancreatic enzymes in the pathogenesis of acute pancreatitis, knowledge of the normal activation process of human zymogens initiated by EP cleaved trypsinogen to trypsin is apparently of great interest. As previous reported, the heavy chain of EP is mandatory for efficient macromolecular substrate recognition ^24^. Subsequently, we tried to capture the substrate engaged EP. Firstly, we incubated the activated mutant EP with its physiological substrate trypsinogen. As validated by distinct migration pattern in native gel electrophoresis, trypsinogen binds firmly to the activated mutant EP (Fig. S8A). Therefore, we further purified this interacting complex by size exclusion chromatography, and chose the right fraction for cryo-EM analysis (fig. S8B, and Table 1). Our initial reconstruction showed obvious orientation preference problem of the particles (Fig. S8C-D). Though tilted 20 degrees for later data acquisition, the orientation preference remains unsolved but partially improved (Fig. S8E-H).

The overall architecture of substrate-bound EP is similar to that of inactive EP structure, exhibiting an extra density attached to CUB2 (Figs. 4A and fig. S9A). This extra density appeared to be able to accommodate trypsinogen, with the model (predicted by AlphaFold2) docked into the density as a rigid body (Figs. 4B and S9B). However, the N-terminal activation peptide need minor manual adjustment to fit properly. This N-terminal tail inserts into the catalytic pocket, near the catalytic triad (Figs. 4C and S9C). Nevertheless, the cleavage site, the specific trypsinogen activated peptide (DDDDK), is ∼20 Å away from the catalytic pocket, indicating this state is not a ready-to-cleave state. This trypsinogen-CUB2 binding is consistent with previous report that CUB2 seems to be essential for recognition of natural substrates ^45, 46^. This discovery solidifies the role of heavy chain in macromolecular substrate recognition and would be the explanation for the slow cleavage of trypsinogen by the isolated light chain ^30^. Noteworthy, MAM had a better resolved density after substrate binding (Fig. 4B). Altogether, this trypsinogen-engaged EP complex provides valuable insights into substrate recognition mechanism.

**Fig. 4.**
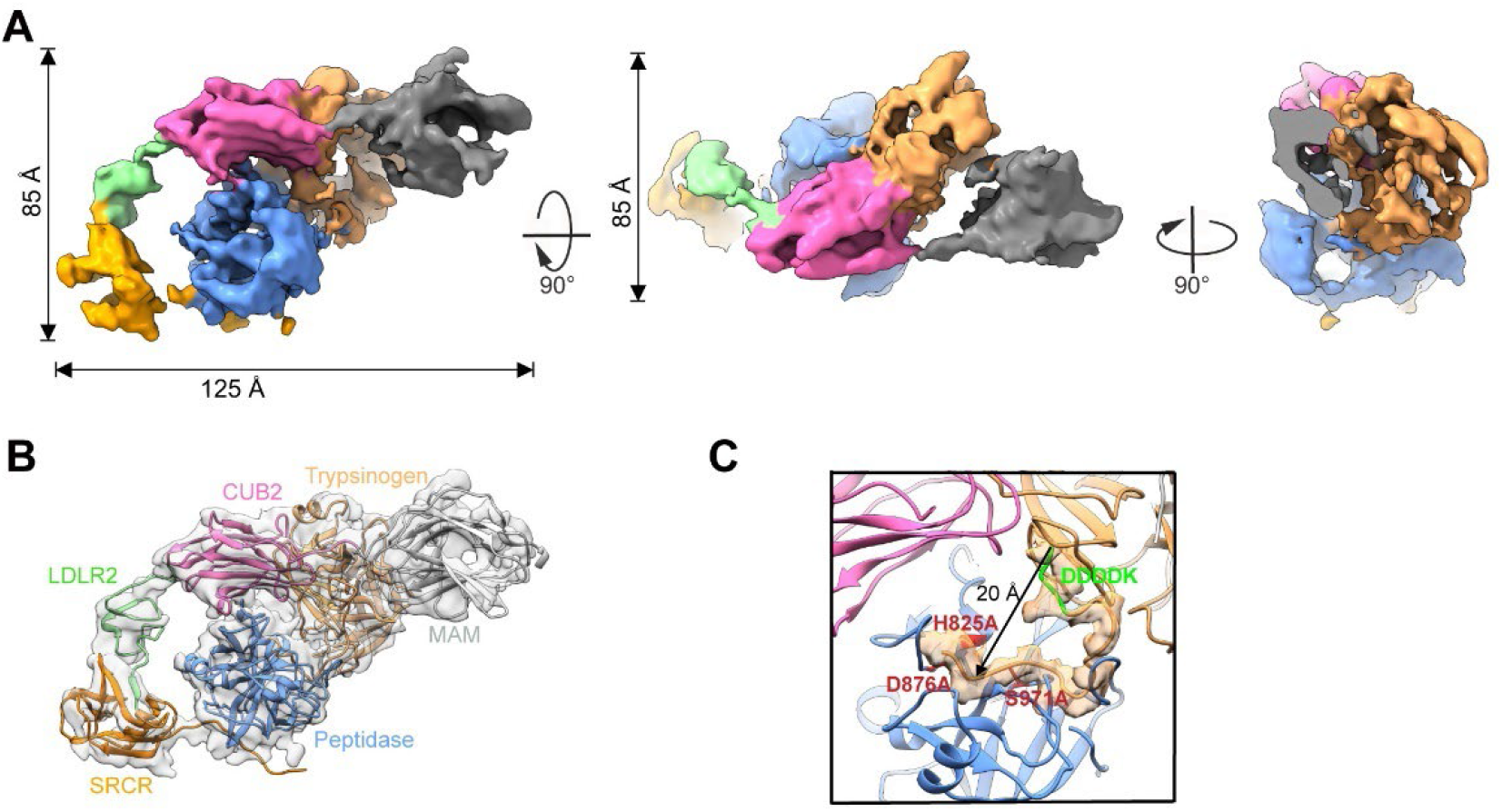
Structural features of substrate-engaged EP. **(A)** Cryo-EM structure of substrate-engaged EP, the extra density corresponding to trypsinogen is colored in sandy brown. **(B)** Model fitting of the substrate-engaged EP map. The additional MAM and trypsinogen were fitted and labeled. **(C)** Substrate enters the catalytic pocket, near the catalytic triad (red). The cleavage site, DDDDK, is also labeled.

## Discussion

Acute pancreatitis is a complex inflammatory disease of the pancreas, with its exact pathogenesis remains unknown ^47, 48^. The inappropriate activation of pancreatic trypsinogen was believed to cause acute pancreatitis, with the underlying molecular mechanism representing a critical clue for pancreatitis therapy ^18^. Acquired EP deficiency may be the origin of indigestion and malabsorption caused by functional pancreatic insufficiency ^13, 19, 49^. EP, the physiological activator of trypsinogen, has been discovered for more than 100 years ^1, 50, 51^. However, the function model for substrate processing by EP remains awaiting the structural evidences. Our structural analysis of EP and its complex with substrate is an important step to decrypt the pathogenesis of pancreatitis and may open a new direction for the treatment of pancreatitis.

We carried out cryo-EM analysis of the EP in inactive, active, inhibited and substrate-bound states and revealed the EP structures with distinct conformations. The inactive EP-core (524-1,019 residues, ∼56KDa) reached 2.7 Å as the highest resolution among all structures, indicating a tight and steady assembly. Our active structures reveal that the trypsin-treated EP remodels the activation domain by the N-terminus of the light chain, turning the whole steady enzyme into the dynamic active conformation. The binding of nafamostat enables us to delineate the most complete profile of EP, representing the general inhibition mechanism by covalent bond and occupancy. While in the substrate-bound structure, the architecture discovered the recognition and binding of macromolecular trypsinogen on CUB2 domain.

Our high-resolution structures detected all the glycan densities (Figs. 1C, 2A, 3D, and S4D), indicating the hEP is in a nearly physiological state. Thus, we proposed a working model for hEP functionality (Fig. 5). It emerged as a zymogen, requiring trypsin or other related proteases to activate ^6, 23, 24^. Initially, the peptidase domain on the light chain was protected by the multi-domains of heavy chain, inaccessible to the substrate. Upon activation, the relative dynamics between the EP-core and the LCM domains might originate from the rapid movement of the core region, with the LCM domains closely connected to the immobilized TM domain. This large flexibility facilitates the substrate recruitment and exposes the catalytic sites for subsequent cleavage. Once trypsinogen bound to CUB2, it reduces the dynamic energy and improves the stability of the complex. After the cleavage of the N-terminal tail, trypsin is released to initiate the pancreatic zymogens cascade. Then the EP goes back to its dynamic state to recruit and catalyze more trypsinogen. These novel discoveries elucidate the activation mechanism of EP toward the catalytic cleavage trypsinogen.

**Fig. 5.**
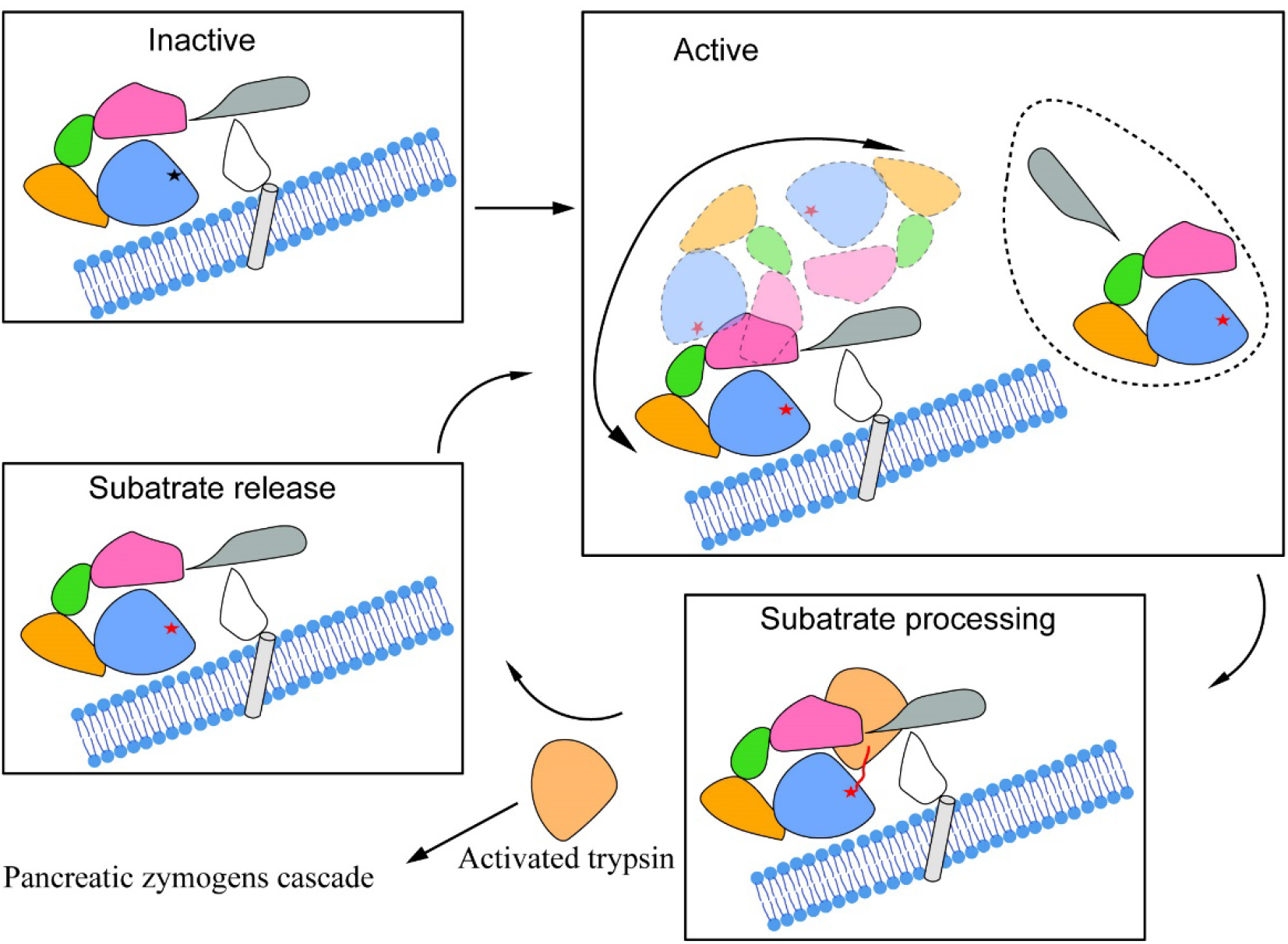
Proposed working model for hEP. In inactive state, the multi-domains of EP heavy chain protect the peptidase domain by surrounding it up. Upon activation, the EP-core become extremely dynamic to facilitate the exposure of the catalytic sites and recruitment of substrates. The complex in the dashed line displays one of the active states stabilized by an inhibitor. Once trypsinogen bound to CUB2, it reduces the dynamic energy and improves the stability of the whole complex. After the cleavage of the N-terminal tail, trypsin released to initiate the pancreatic zymogens cascade. And the EP goes back to its dynamic state to find and catalyze more substrates.

Noteworthy, our inhibitor assays screened a few small molecules, and nafamostat was found to be the most effective candidate to inhibit EP. Furthermore, the covalent binding of nafamostat was identified within the catalytic pocket of the high-resolution EP complex structure. Nafamostat would potentially serve as a drug candidate to target EP-redundant disease, besides the usage of anti-coagulation therapy in acute pancreatitis.

In summary, we solved a series of hEP structures, including the different states of its functional cycle. The work delineates the structural basis for hEP activation and substrate recognition, establishing a foundation for the study of hEP structure-function relationships. These mechanism discoveries and inhibition assays shed light on nafamostat-derived drug research and development, having profound pathobiological and therapeutic targeting implications.

## Methods

### Protein expression and purification

The wild type hEP (182-1,019aa) and mutant hEP (H825A, D876A and S971A, 182-1,019aa) were cloned with a N-terminal signal peptide and C-terminal 10×His tags using homologous recombination (HieffCloneTM One Step Cloning Kit, Yeasen). The plasmids were transient transfected into HEK293F, which further cultured at 37°C for 6 days. Cells were harvested, suspended, and lysed using a French press. The cellular debris were cleared by centrifugation and the supernatant was loaded onto a Ni-NTA affinity purification column. The eluted fractions containing hEP was detected using SDS-PAGE, pooled and dialyzed into buffer (20 mM Tris-HCl, 20 mM NaCl, pH 7.6). The final proteins were checked by SDS-PAGE and native PAGE. For the native PAGE, the samples were mixed with native dye and subjected to 8% native-PAGE. We ran the gel at 120 V at 4 °C for 1 h to separate EP in different conditions. The gels were stained by coomassie blue.

### Preparation of activated hEP by Trypsin

Trypsin (T1426, Sigma-Aldrich) was incubated with purified hEP in buffer (20mM Tris-HCl, pH7.6, 150mM NaCl, and 5mM CaCl2) at a final mass ratio of 1: 100 for 2hrs at 37°C. The activity was determined by using the enteropeptidase activity assay kit (K758-100, Biovision). The final proteins were checked by SDS-PAGE and native PAGE.

### Preparation of inhibited hEP

Different inhibitors were tested first to find the best inhibitor of hEP, including nafamostat, camostat, gabexate, PMSF, and ulinastatin. The inhibitors were incubated with the activated hEP for 30min at 37°C prior the EP activity assay.

To find the best inhibit efficacy of nafamostat (10 mg/ml), several concentrations were tested, with the molar ratio to activated hEP at 50:1, 100:1, 500:1, and 1000:1. Inhibitor solutions were transferred to assay the EP activity using enteropeptidase activity assay kit (K758-100, Biovision).

### Preparation of substrate-bound hEP

A 1:5 molar ratio of the mutated hEP to trypsinogen was incubated in PBS at 37°C for 30min, following the detection of EP catalytic ability by using the trypsin activity kit (MAK290, Sigma-Aldrich), and cross-linking by 0.05% glutaraldehyde for complex stabilization. The cross-linking was stopped after 30min incubation at room temperature by adding of Tris, pH 8.0, to a final concentration of 1 mM. To obtain the highest purity, gel filtration was performed in 20mM Tris-HCl, pH7.6, 150mM NaCl, and 5mM CaCl2. The fractions containing trypsinogen-hEP was detected using SDS-PAGE and native PAGE, and proteins were concentrated for cryo-EM data collection.

### Surface plasmon resonance analysis

Commercial EP was immobilized onto a CM5 sensor chip surface with the NHS/EDC method using Biacore 8k (Cytiva) and 1× PBS-T running buffer (1× PBS with 0.05% Tween-20). The nafamostat flew over the chip at a rate of 30 μl /min. After saturated with the nafamostat, the TMPRSS15 was injected at the same rate for another 120s. The data was fitted by using Affinity (Cytiva).

### Cryo-EM sample preparation and data collection

3 μl samples at 1-1.5mg/ml were placed onto a glow-discharged holey amorphous nickel-titanium alloy film supported by 400 mesh gold grids ^52^, followed by blotting utilizing Vitrobot Mark IV (FEI/Thermo Fisher Scientific), and flash frozen in liquid ethane.

Images were taken by using a Titan Krios transmission electron microscope (ThermoFisher) operated at 300 kV, equipped with a Bio Quantum post-column energy filter with zero-loss energy selection slit set to 20 eV. Images were collected by using a K2 Summit direct electron detector (Gatan) in super-resolution counting mode, corresponding to a pixel size of 0.523 Å at the specimen level. Each movie was dose-fractioned into 36 frames with a dose rate of 8 e^-^ per pixel per sec on the detector. The exposure time was 7.2 s with 0.2 s for each frame, generating a total dose of ∼ 52 e^-^/Å^2^. Defocus values varied from -1.5 to -2.5 μm. All of the images were collected by utilizing the SerialEM automated data collection software package ^53^.

### Cryo-EM data processing

Unless otherwise specified, single-particle analysis was mainly executed in cryoSPARC ^54^, including patch motion correction, patch CTF estimation. For the inactive dataset, the initial particles were picked by Blob picker. Good templates were selected by confirming the general features of EP through 2D and 3D classifications (particles binned 4 × 4 during extraction), which used for template-based particle picking of all the datasets. Four rounds of heterogeneous refinement were performed to classify the particles from template picker, resulting one class with good features. The selected good classes from Blob and template pickers were combined together, further classified by two rounds of heterogeneous refinement. The particles from the good class were re-extracted with original size (pixel size=1.046 Å), and were subjected to further classification. The bad class was removed to obtain a cleaner particle stack, the remaining three classes were combined to reconstructed into a complete map, while the core map were local refined by applying a core mask. The locally refined EP-core and the low-resolution region in the whole map were merged in UCSF Chimera and used for subsequent model building and analysis.

For all the other datasets, the selected reference and good particles from the inactive dataset performed the template for particle picking and seed for seed-facilitated 3D classification ^55^. Directional Fourier Shell Correlation (dFSC) curves are calculated as described ^56^, and the nominal resolution is estimated from the averaged FSC using FSC = 0.143 criterion ^57^. The local resolution was estimated either by using ResMap ^58^ or deepEMhancer ^59^.

### Pseudo-atomic model building and validation

The inactive model of hEP predicted by AlphaFold2 ^60, 61^ was initially fitted in the density maps with UCSF Chimera ^62^ and manually adjusted in COOT ^63^. All models were refined against the corresponding maps multiple rounds using real-space refinement in PHENIX ^64^, where the last iteration included ADP refinement to calculate the B-factors. The nafamostat and glycans were added in COOT ^63^.

Model validation was performed following the protocol of phenix.molprobity ^64, 65^. UCSF Chimera ^62^ and ChimeraX ^66^ was used for map segmentation and figure generation.

## Acknowledgements

We would like to thank the cryo-EM facility of Southern University of Science and Technology for providing facility support. We thank Hong Wu and Zhangyun Jiang from Shanghai YueXin Life-Science Information Technology Co., Ltd for help with computation. This work was supported by National Natural Science Foundation of China Grants 82020108005 (to Z. L.) and 82022008 (to H. H.); and grant from technology support plan of China 2015BAI13B08 (to Z. L.).

## Author contributions

Conceived and designed the experiments: X.Y., Z.D., L.P., and H.H. Designed constructs for structural studies: X.Y., L.P. and Q.S. Protein purification: L.P., Q.S., F.C., and D.Z. Performed functional analysis: X.Y., Z.D., C.X., K.L., H.Y., S.L., Z.L., and H.H. Performed EM data collection and analysis: X.Y., and Z.D. Structure reconstruction: X.Y., and Z.D. Model building: Z.D. SPR analysis: X.Y. and L.P. Wrote the paper: X.Y., Z.D., L.P., and H.H.

## Accession numbers

EM maps have been deposited in the Electron Microscopy Data Bank under accession codes of EMD-32715, EMD-32714, EMD-32716, EMD-32717, EMD-32828, and EMD-32829 for EP in inactive, active-wt, active-mut, inhibited-core, inhibited-complete, and substrate-bound states, respectively. Pseudo-atomic models have been deposited in the Protein Data Bank under accession numbers of 7WQX, 7WQW, 7WQZ, and 7WR7 for EP in inactive, active-wt, active-mut, and inhibited states, respectively.

## Figures and legends

**Fig. S1.**
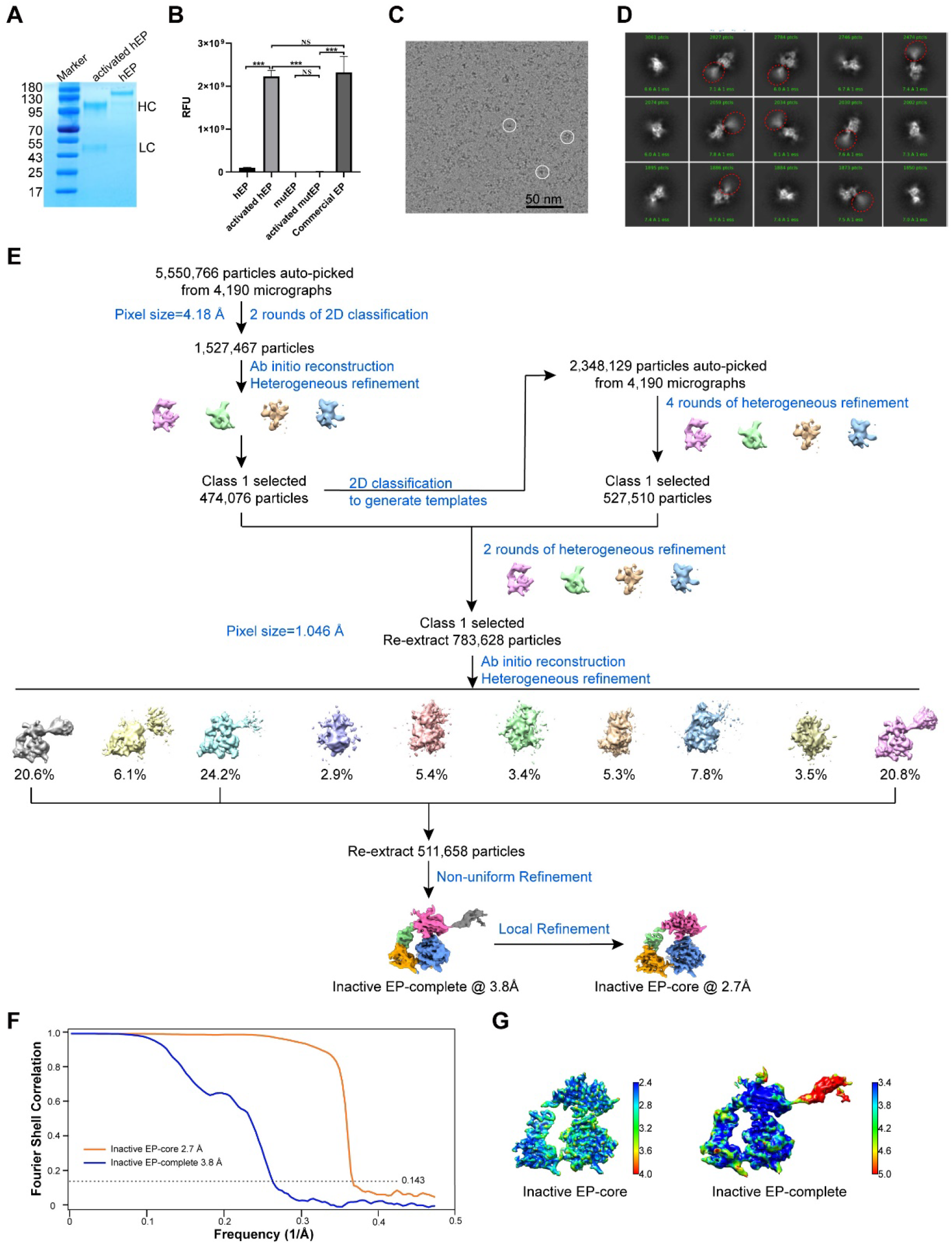
Biochemical and cryo-EM analysis of hEP in the inactive state. **(A)** Identification of hEP by SDS-PAGE, followed by coomassie brilliant blue staining. EP activation is realized by trypsin cleavage into heavy chain (HC) and light chain (LC). **(B)** *In vitro* EP activity experiment. SDs (standard deviations) were calculated from at least three independent experiments. Significance was tested using one-way ANOVA; ^***^*P*<0.001, ^NS^*P*>0.05. **(C)** Representative cryo-EM micrograph of the inactive hEP. For better visualization, the original data was low-pass filtered to enhance the contrast. The scale bar indicates 50 nm. (**D)** Representative reference-free 2D class averages for inactive EP particles. The fuzzy domains were circled by red ellipses. (**E)** Work flow for the processing of inactive EP data. **(F)** Resolution evaluation of the cryo-EM maps by Fourier shell correlation (FSC) 0.143 criterion. **(G)** Local resolutions of the cryo-EM maps determined using ResMap, with the color bar on the right labeling the resolutions (in Å).

**Fig. S2.**
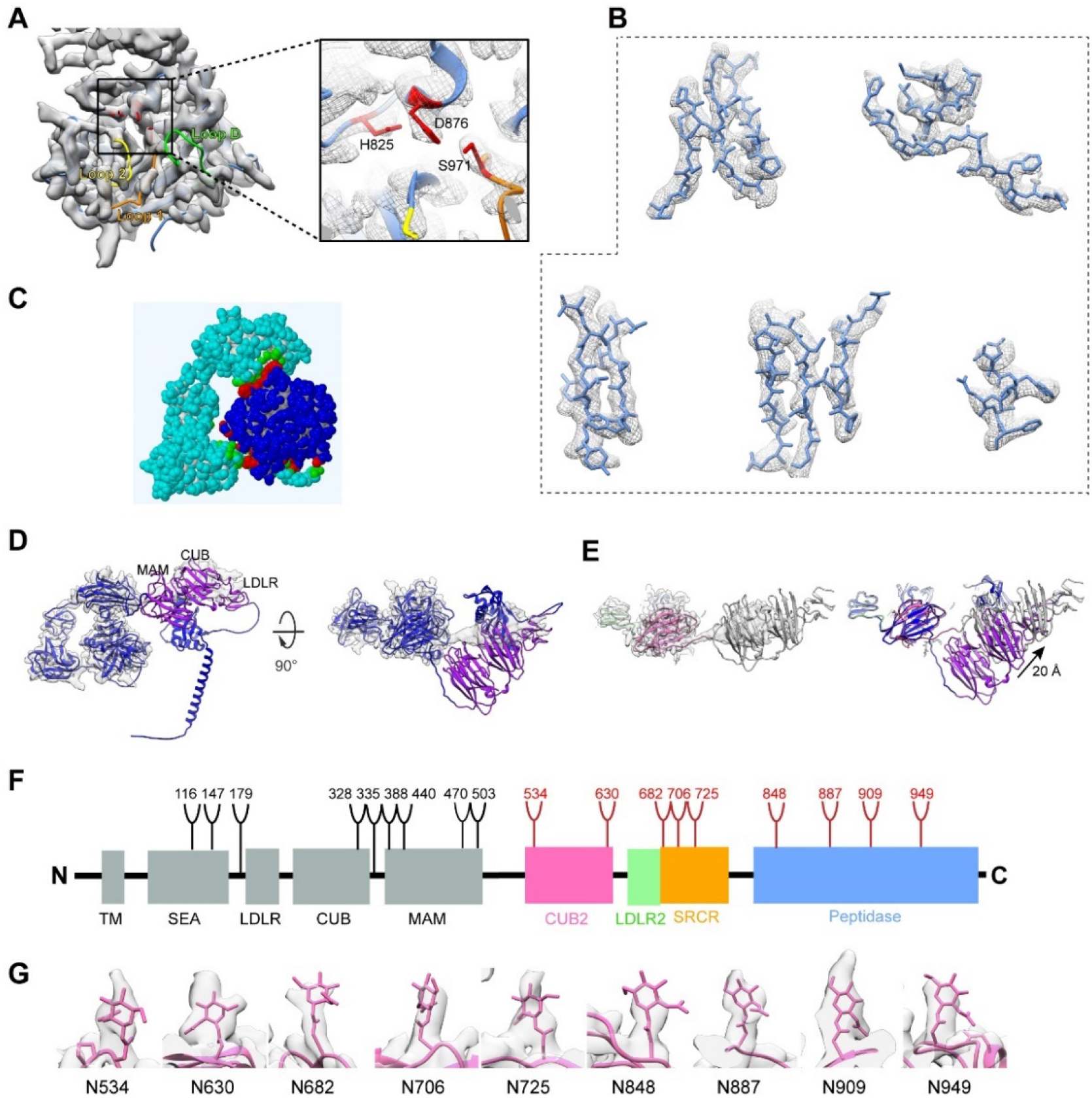
Structural features of hEP in the inactive state. **(A)** Inactive EP map fitted with the corresponding pseudo-atomic model, with the catalytic triad colored in red and enlarged for clarity. The surface loops, L1, L2, and LD, are indicated. **(B)** The good fitting of β-strand and α-helix showing high resolution features. **(C)** The interaction mode between heavy and light chain, with the interacting residues colored in red. **(D)** The AF2 predicted full-length EP model fits in our inactive EP map. **(E)** Our EP-complete model fits in complete map. The overlaying of the model (grey) with the AF2 predicted model (with color) shows a slightly shift. **(F)** Schematic representation of N-linked glycans in EP. The positions of N-linked glycosylation sequins are shown as branches. 9 N-linked glycans detected in our cryo-EM are shown in red, the remaining undetected ones in black. **(G)** The good fitting of 9 N-linked glycans detected in our inactive EP.

**Fig. S3.**
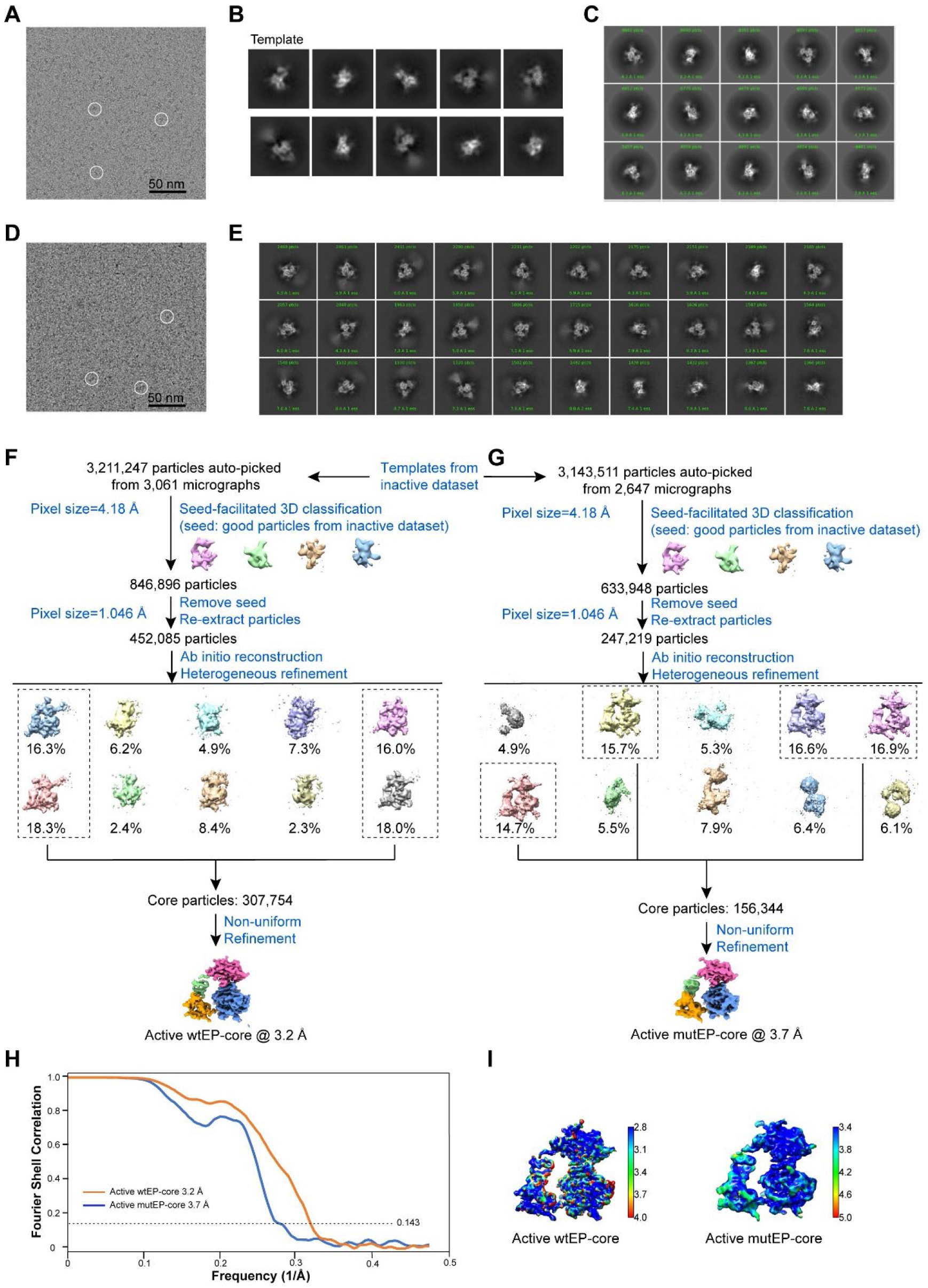
Cryo-EM analysis of hEP in the active state. **(A)** Representative cryo-EM micrograph of the active wtEP. For better visualization, the original data was low-pass filtered to enhance the contrast. The scale bar indicates 50 nm. (**B)** Class-averages from the inactive dataset serving as the templates for autopicking. **(C)** Representative reference-free 2D class averages for active wtEP-core particles. **(D)** Representative cryo-EM micrograph of the active mutEP. For better visualization, the original data was low-pass filtered to enhance the contrast. The scale bar indicates 50 nm. (**E)** Representative reference-free 2D class averages for active mutEP particles. (**F-G)** Work flows for the processing of active wtEP **(F)** and mutEP **(G)** data. **(H)** Resolution evaluation of the cryo-EM maps by Fourier shell correlation (FSC) 0.143 criterion. **(I)** Local resolutions of the cryo-EM maps determined using ResMap, with the color bar on the right labeling the resolutions (in Å).

**Fig. S4.**
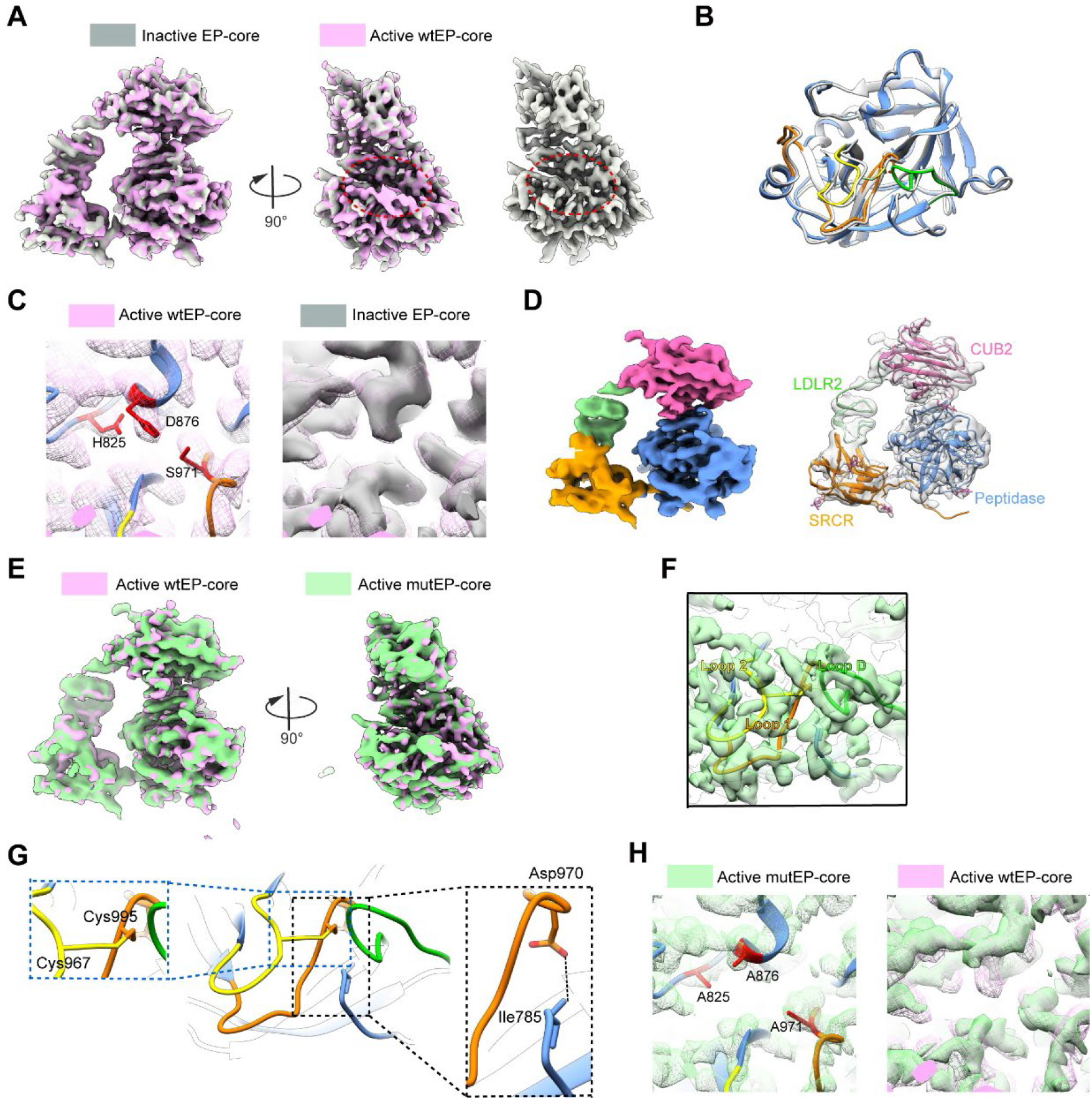
Structural features of hEP in the active state. **(A)** Overlay of inactive EP-core (grey) with active EP-core (pink) maps, and the extra density near the active pocket were circled by a dashed red ellipse. **(B)** Structural overlaying between our structure (with different colors) and published model 4DGJ (grey) shows little variations. **(C)** The fitting of the catalytic triad in active wtEP-core map and the overlaying of this region with that in the inactive EP-core map. **(D)** Cryo-EM structure of active mutEP-core, with model fitted in. The detected 9 N-linked glycans were indicated by atomic view. **(E)** Overlay of active wtEP-core (pink) with active mutEP-core (light green) maps. **(F)** The fitting of the surface loops in active mutEP-core map. **(G)** The activation domain in active mutEP-core structure. The interdomain disulfide pair (Cys967-Cys995) and the Asp970:Ile785 salt bridge were enlarged for clarity. **(H)** The fitting of the mutated catalytic triad in active mutEP-core map and the overlaying of this region with that in the active wtEP-core map.

**Fig. S5.**
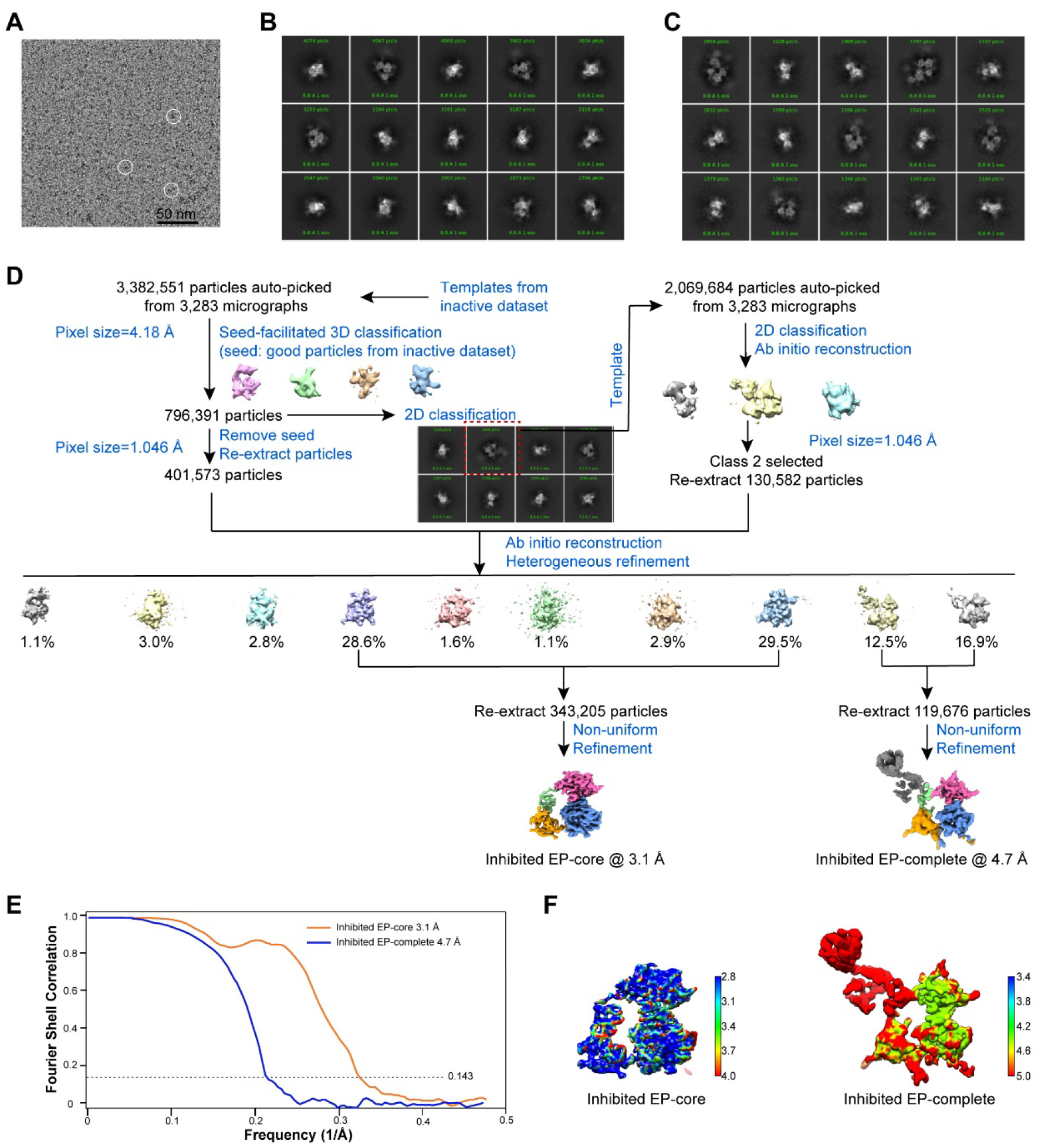
Cryo-EM analysis of hEP in the inhibited state. **(A)** Representative cryo-EM micrograph of inhibited hEP. For better visualization, the original data was low-pass filtered to enhance the contrast. The scale bar indicates 50 nm. (**B)** Representative reference-free 2D class averages for inhibited EP-core particles. **(C)** Representative reference-free 2D class averages for inhibited EP-complete particles. (**D)** Work flow for the processing of inhibited hEP data. **(E)** Resolution evaluation of the cryo-EM maps by Fourier shell correlation (FSC) 0.143 criterion. **(F)** Local resolutions of the cryo-EM maps determined using ResMap, with the color bar on the right labeling the resolutions (in Å).

**Fig. S6.**
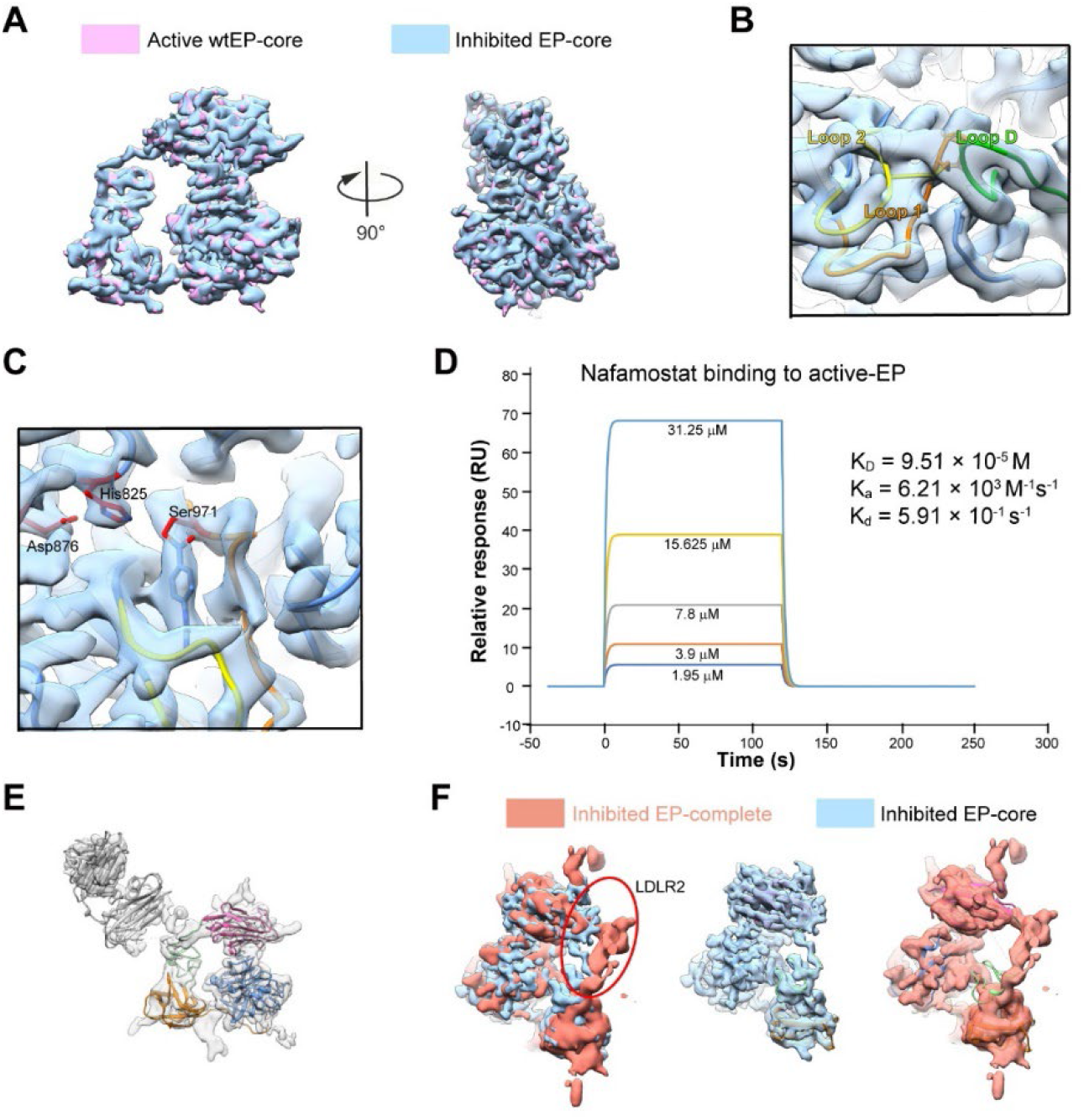
Structural features of hEP in the inhibited state. **(A)** Overlay of inhibited EP-core (blue) with active wtEP-core (pink) maps. **(B)** The fitting of the surface loops in inhibited EP-core map. **(C)** Interactions between nafamostat and hEP. **(D)** Surface plasmon resonance analysis of hEP with nafamostat. **(E)** The structure model of the inhibited EP-complete, with the map in transparent. **(F)** The conformational change of LDLR2 due to the relocation of the LCM domains.

**Fig. S7.**
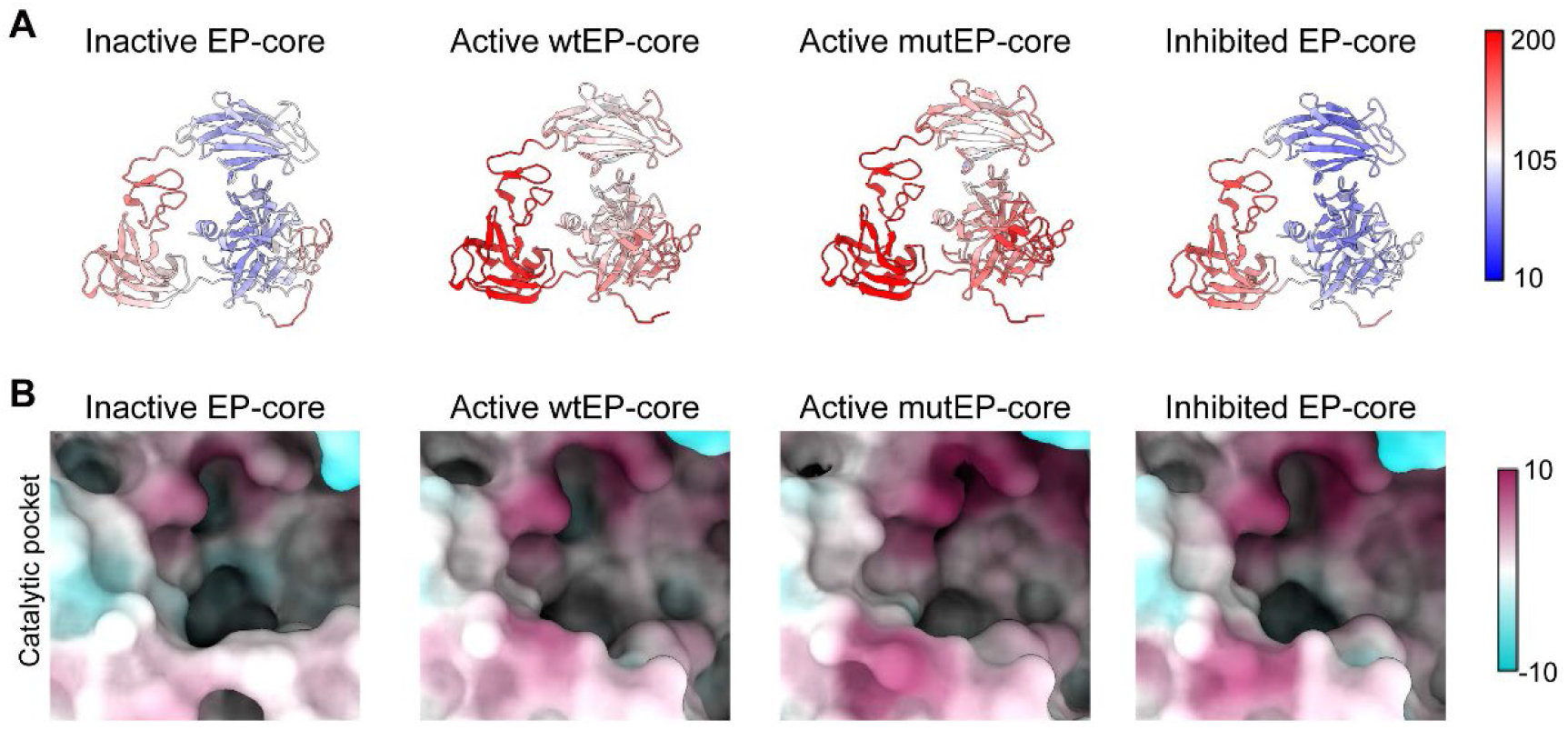
A comparison of hEP conformations in different states. **(A)** Atomic models colored according to the B-factor distribution (ranging from 10 Å^2^ (blue) to 200 Å^2^ (red)). **(B)** Surface property of hEP, with the most distinct residues in catalytic pocket region indicated.

**Fig. S8.**
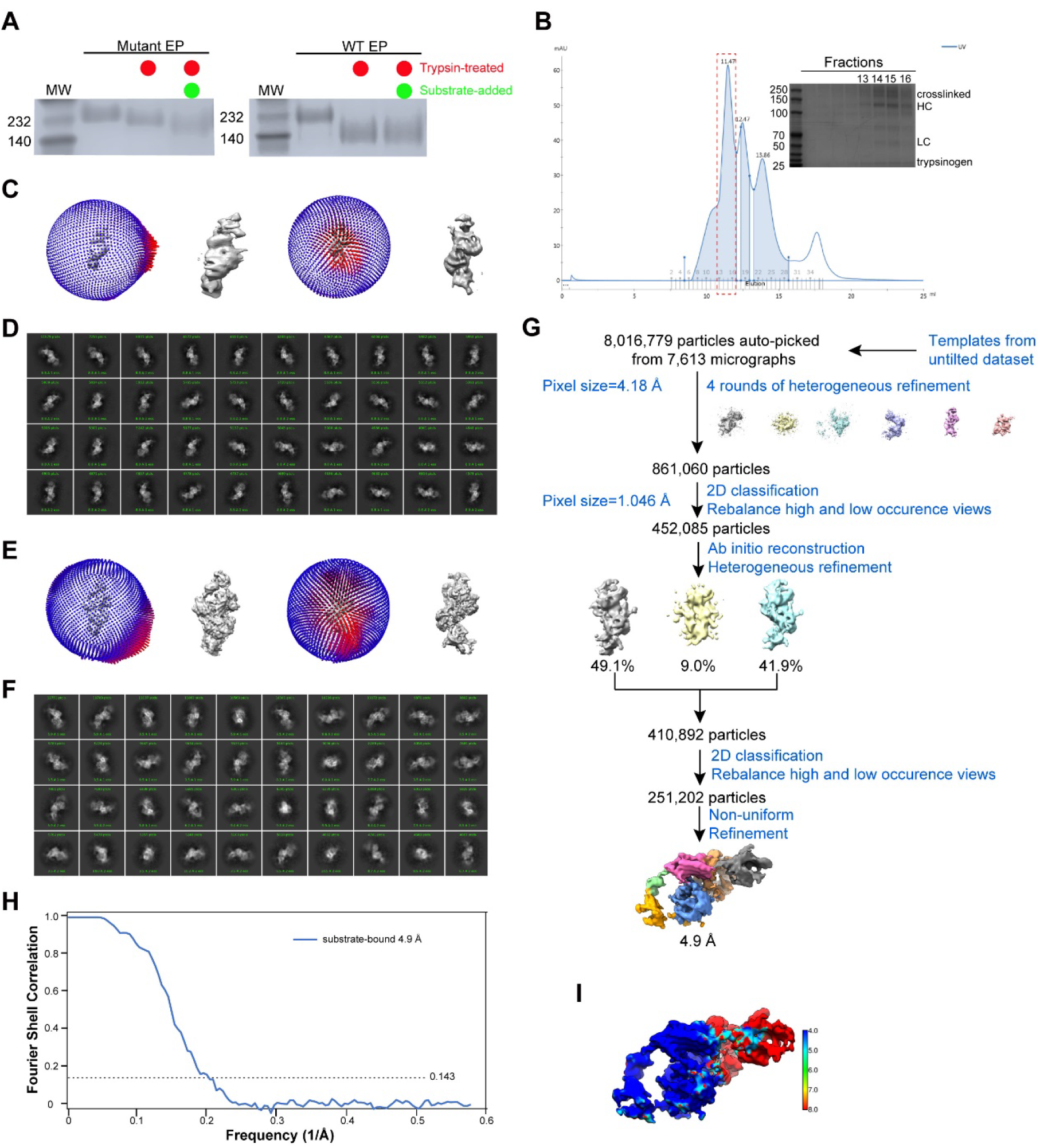
Cryo-EM analysis of hEP in the substrate-engaged state. **(A)** The association of trypsinogen to the mutate EP monitored by a gel-shift assay. In the presence of substrate, the mutEP shows a distinct migration pattern, while no observable changes for the wtEP. **(B)** Different compositions of EP-trypsinogen were fractionated by size exclusion chromatography. Peak fractions were analyzed by SDS-PAGE. **(C)** The angle distributions of the substrate-engaged data without tilting. **(D)** The representative reference-free 2D class averages for the substrate-engaged data without tilting. **(E)** The angle distributions of the substrate-engaged data with 20 degrees tilting. **(F)** The representative reference-free 2D class averages for the substrate-engaged data with 20 degrees tilting. (**G)** Work flow for the processing of substrate-engaged data. **(H)** Resolution evaluation of the cryo-EM maps by Fourier shell correlation (FSC) 0.143 criterion. **(I)** Local resolutions of the cryo-EM map determined using ResMap, with the color bar on the right labeling the resolutions (in Å).

**Fig. S9.**
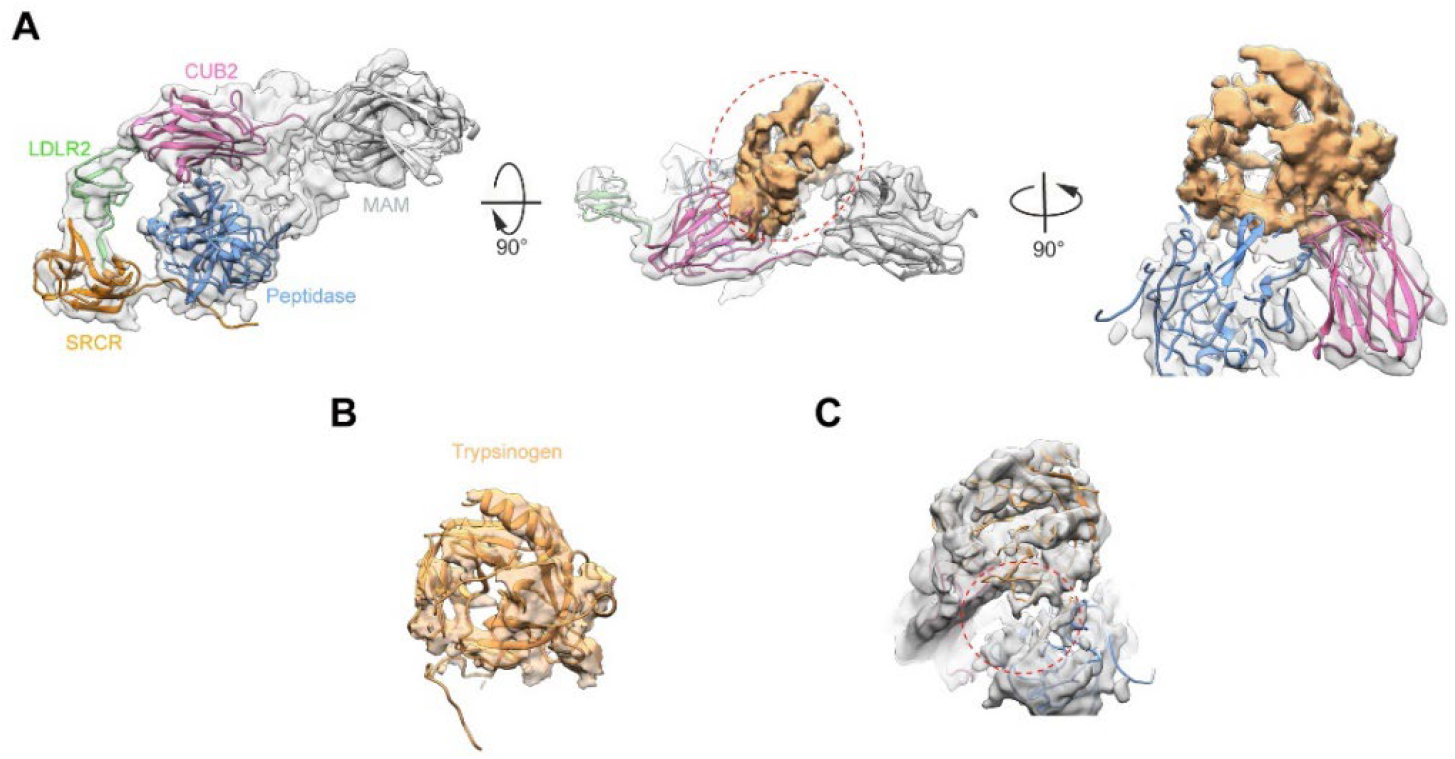
Validation of the extra density to be trypsinogen in substrate-engaged map. **(A)** Model fitting of the substrate-engaged EP map showing the location of the extra density. The density was subtracted and colored in sandy brown. **(B)** Fitting of the trypsinogen model in the subtracted extra density. **(C)** Substrate enters the catalytic pocket, indicated by a red ellipse.

**Table S1.** Cryo-EM data collection, reconstruction, and refinement statistics.

|  | Inactive-EP | Active wtEP-core | Active mutEP-core | Inhibited EP-core | Inhibited EP-complete | Substrate-bound |
| --- | --- | --- | --- | --- | --- | --- |
| **PDB** | 7WQX | 7WQW | 7WQZ | 7WR7 | N/A | N/A |
| **EMDB** | 32715 | 32714 | 32716 | 32717 | 32828 | 32829 |
| **Data collection** |  |  |  |  |  |  |
| EM equipment | FEI Titan Krios | FEI Titan Krios | FEI Titan Krios | FEI Titan Krios | FEI Titan Krios | FEI Titan Krios |
| Voltage, kV | 300 | 300 | 300 | 300 | 300 | 300 |
| Detector | K2 | K2 | K2 | K2 | K2 | K3 |
| Pixel size, Å | 0.523 | 0.523 | 0.523 | 0.523 | 0.523 | 0.425 |
| Electron dose, e/Å<sup>2</sup> | 52 | 52 | 52 | 52 | 52 | 50 |
| Exposure time, s | 7.2 | 7.2 | 7.2 | 7.2 | 7.2 | 2.18 |
| Frames | 36 | 36 | 36 | 36 | 36 | 40 |
| Dedocus range, µm | 1.0-3.5 | 0.7-3.4 | 0.9-3.5 | 1.0-3.5 | 1.0-3.5 | 1.4-2.4 |
| **Reconstruction** |  |  |  |  |  |  |
| Software | cryosparc | cryosparc | cryosparc | cryosparc | cryosparc | cryosparc |
| Raw micrographs | 4,190 | 3,061 | 2,647 | 3,283 | 3,283 | 7,613 |
| Final particles | 511,658 | 307,754 | 156,344 | 343,205 | 119,676 | 251,202 |
| Symmetry | C1 | C1 | C1 | C1 | C1 | C1 |
| Final resolution, Å | 2.7/3.8 | 3.2 | 3.7 | 3.1 | 4.7 | 4.9 |
| Map-sharpening B factor, Å<sup>2</sup> | -87.7/-171.7 | -99.7 | -112.4 | -110.9 | -140.0 | -304.7 |
| **Refinement** |  |  |  |  |  |  |
| Software | phenix | phenix | phenix | phenix | N/A | N/A |
| Rms deviations |  |  |  |  |  |  |
| Bond length, Å | 0.0020 | 0.0015 | 0.0017 | 0.0018 | N/A | N/A |
| Bond angle, ° | 0.47 | 0.42 | 0.49 | 0.48 | N/A | N/A |
| Ramachandran plot statistics, % |  |  |  |  |  |  |
| Preferred | 93.99 | 93.29 | 91.46 | 91.26 | N/A | N/A |
| Allowed | 6.01 | 6.71 | 8.54 | 8.74 | N/A | N/A |
| Outlier | 0.00 | 0.00 | 0.00 | 0.00 | N/A | N/A |

